## Supplementary Information for "MetaScore: A novel machine-learning based approach to improve traditional scoring functions for scoring protein-protein docking conformations"

Supplementary Table S1. BLAST parameters.

| Query Length | Substitution Matrix | Gap Open Cost | Gap Extend Cost |
| --- | --- | --- | --- |
| <35 | PAM-30 | 9 | 1 |
| 35-50 | PAM-70 | 10 | 1 |
| 50-85 | BLOSUM-80 | 10 | 1 |
| >85 | BLOSUM-62 | 11 | 1 |

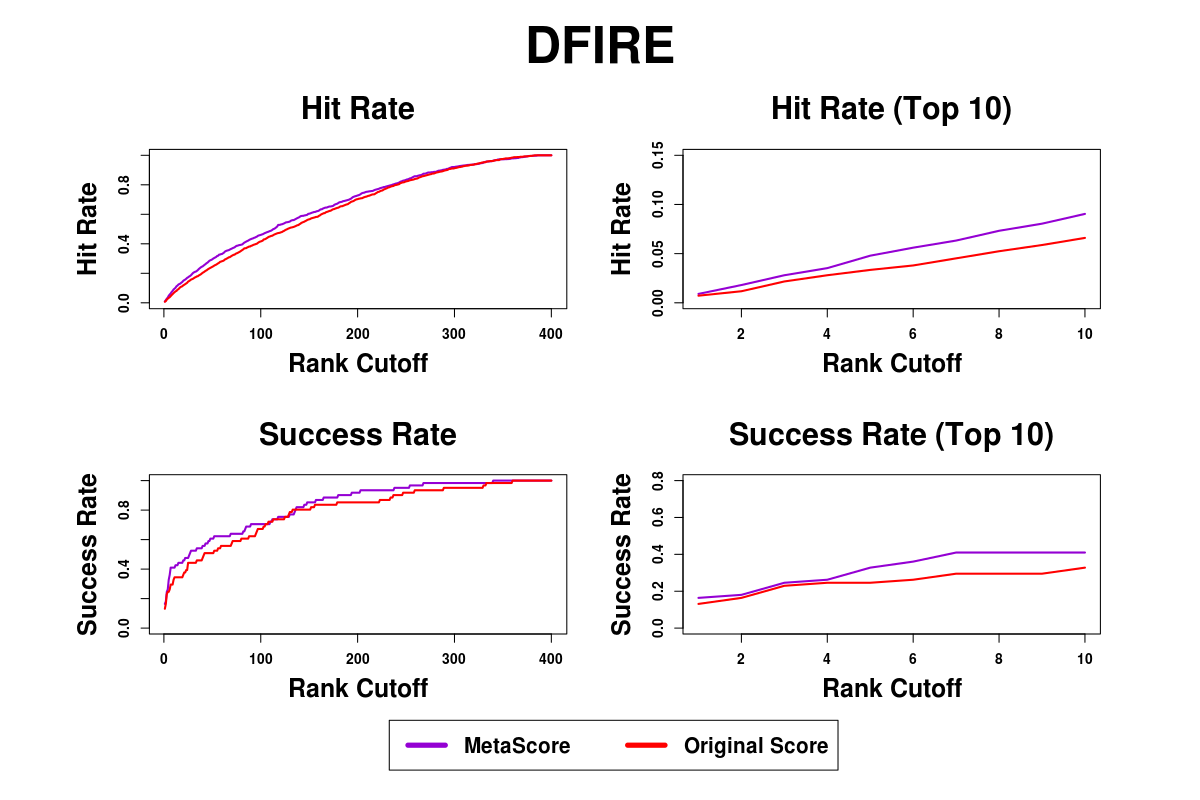

Supplementary Figure S1. Success rates and hit rates plotted against the top m conformations for a classical scoring method (DFIRE), machine learning-based method using RF, and the combined method of the two methods using BM4 decoy set. There are four panels. Top-left panel shows hit rates for conformations of top m ranging from 1 to 400; Top-right panel shows hit rates for conformations of top m ranging from 1 to 10; Bottom-left panel shows success rates for conformations of top m ranging from 1 to 400; Bottom-right panel shows success rates for conformations of top m ranging from 1 to 10.

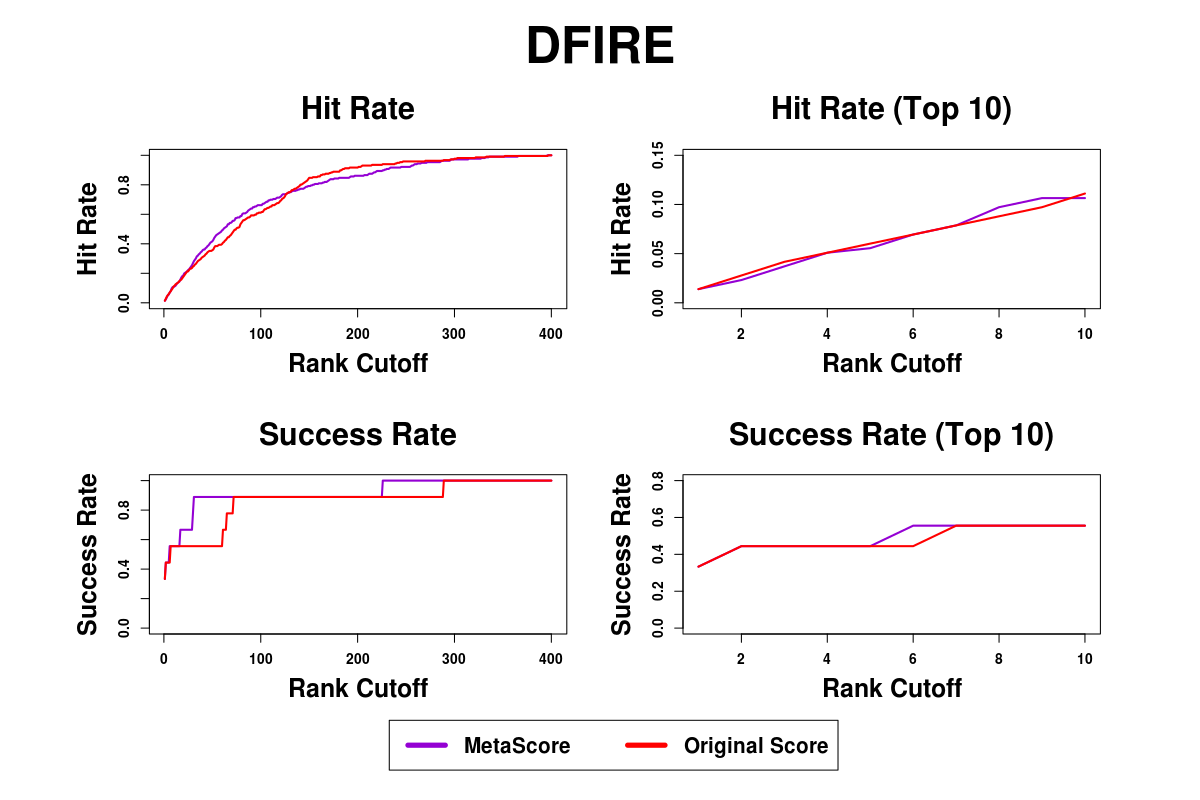

Supplementary Figure S2. Success rates and hit rates plotted against the top m conformations for a classical scoring method (DFIRE), machine learning-based method using RF, and the combined method of the two methods using BM5 decoy set. There are four panels. Top-left panel shows hit rates for conformations of top m ranging from 1 to 400; Top-right panel shows hit rates for conformations of top m ranging from 1 to 10; Bottom-left panel shows success rates for conformations of top m ranging from 1 to 400; Bottom-right panel shows success rates for conformations of top m ranging from 1 to 10.

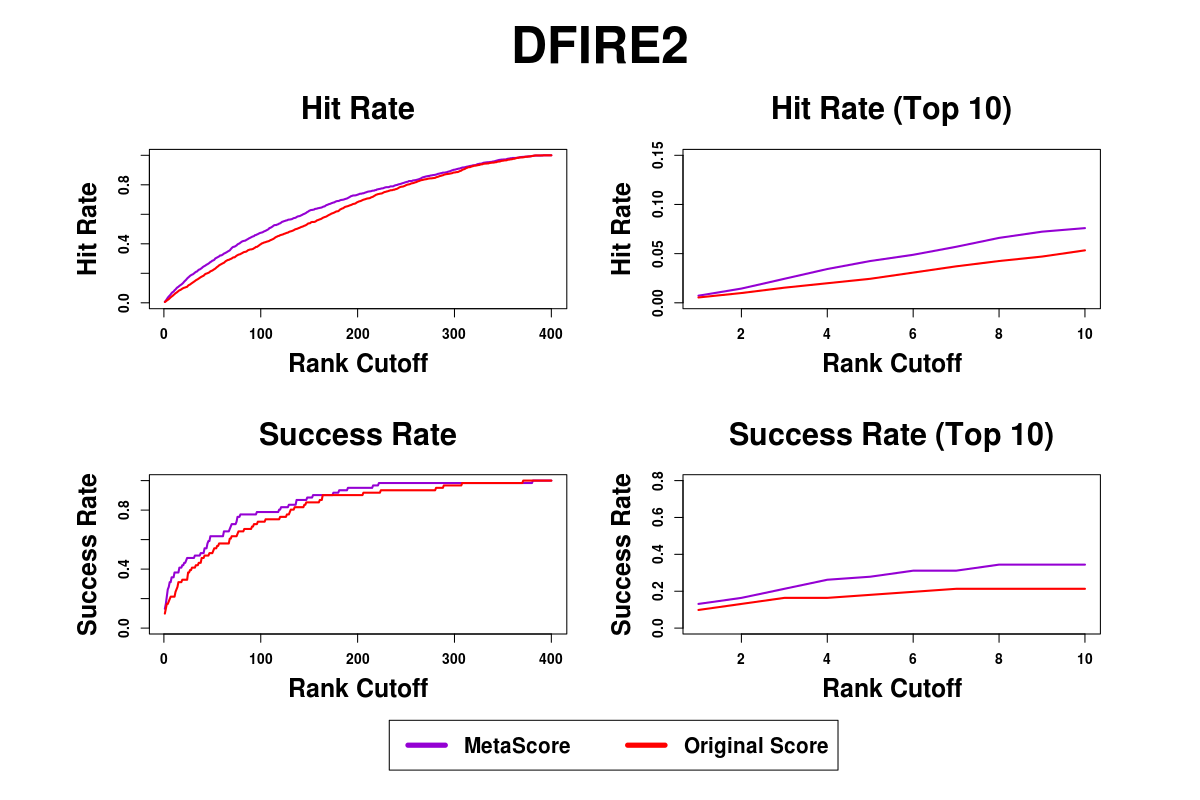

Supplementary Figure S3. Success rates and hit rates plotted against the top m conformations for a classical scoring method (DFIRE2), machine learning-based method using RF, and the combined method of the two methods using BM4 decoy set. There are four panels. Top-left panel shows hit rates for conformations of top m ranging from 1 to 400; Top-right panel shows hit rates for conformations of top m ranging from 1 to 10; Bottom-left panel shows success rates for conformations of top m ranging from 1 to 400; Bottom-right panel shows success rates for conformations of top m ranging from 1 to 10.

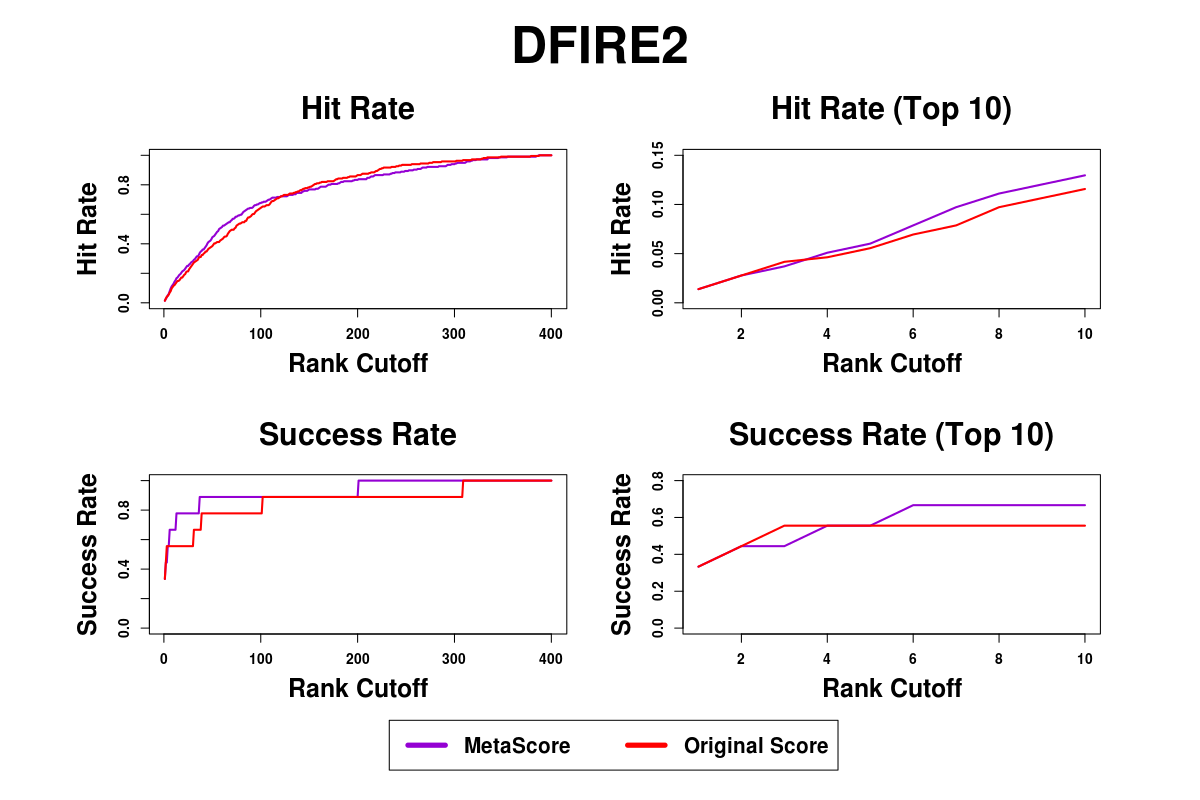

Supplementary Figure S4. Success rates and hit rates plotted against the top m conformations for a classical scoring method (DFIRE2), machine learning-based method using RF, and the combined method of the two methods using BM5 decoy set. There are four panels. Top-left panel shows hit rates for conformations of top m ranging from 1 to 400; Top-right panel shows hit rates for conformations of top m ranging from 1 to 10; Bottom-left panel shows success rates for conformations of top m ranging from 1 to 400; Bottom-right panel shows success rates for conformations of top m ranging from 1 to 10.

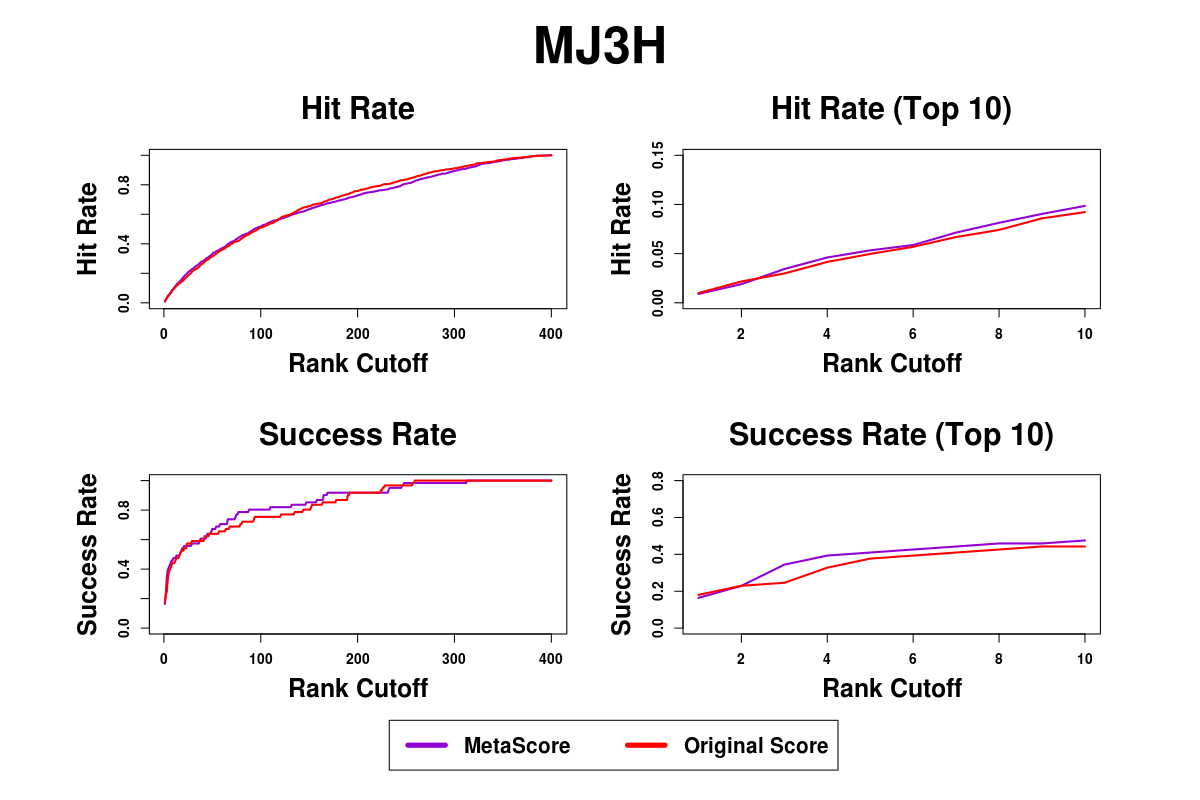

Supplementary Figure S5. Success rates and hit rates plotted against the top m conformations for a classical scoring method (MJ3H), machine learning-based method using RF, and the combined method of the two methods using BM4 decoy set. There are four panels. Top-left panel shows hit rates for conformations of top m ranging from 1 to 400; Top-right panel shows hit rates for conformations of top m ranging from 1 to 10; Bottom-left panel shows success rates for conformations of top m ranging from 1 to 400; Bottom-right panel shows success rates for conformations of top m ranging from 1 to 10.

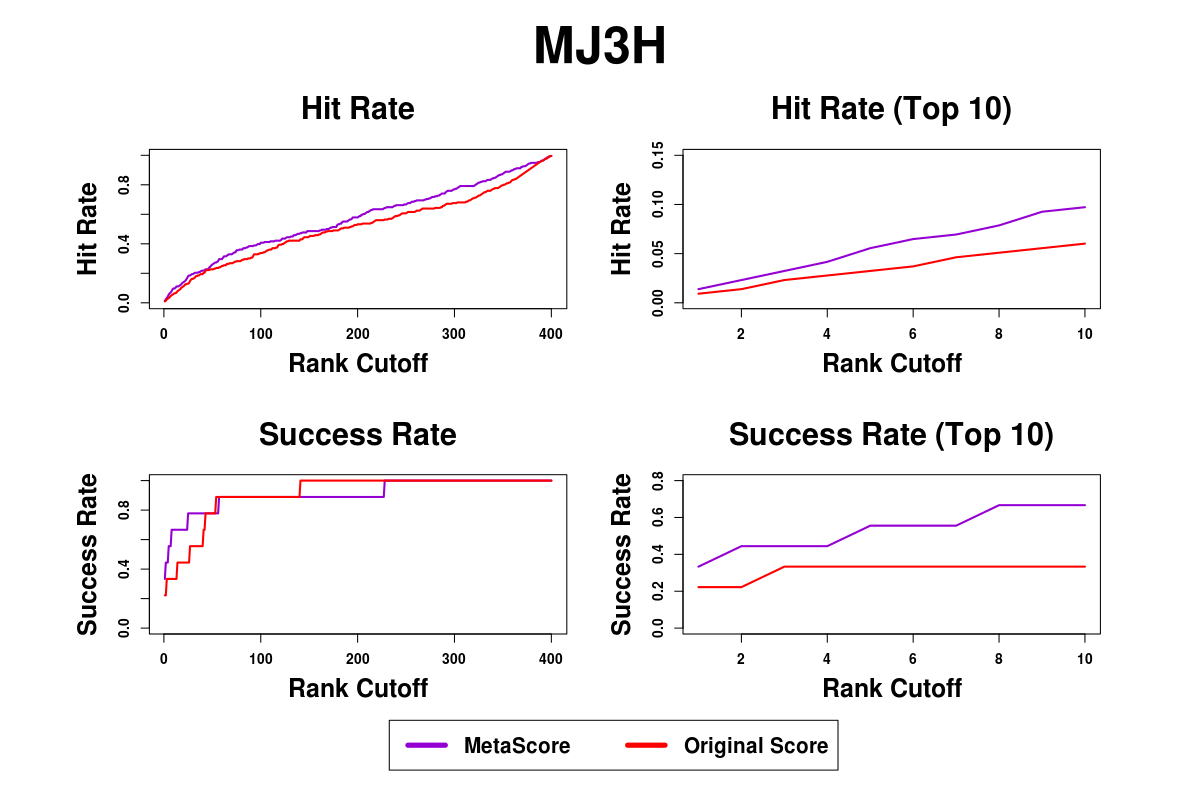

Supplementary Figure S6. Success rates and hit rates plotted against the top m conformations for a classical scoring method (MJ3H), machine learning-based method using RF, and the combined method of the two methods using BM5 decoy set. There are four panels. Top-left panel shows hit rates for conformations of top m ranging from 1 to 400; Top-right panel shows hit rates for conformations of top m ranging from 1 to 10; Bottom-left panel shows success rates for conformations of top m ranging from 1 to 400; Bottom-right panel shows success rates for conformations of top m ranging from 1 to 10.

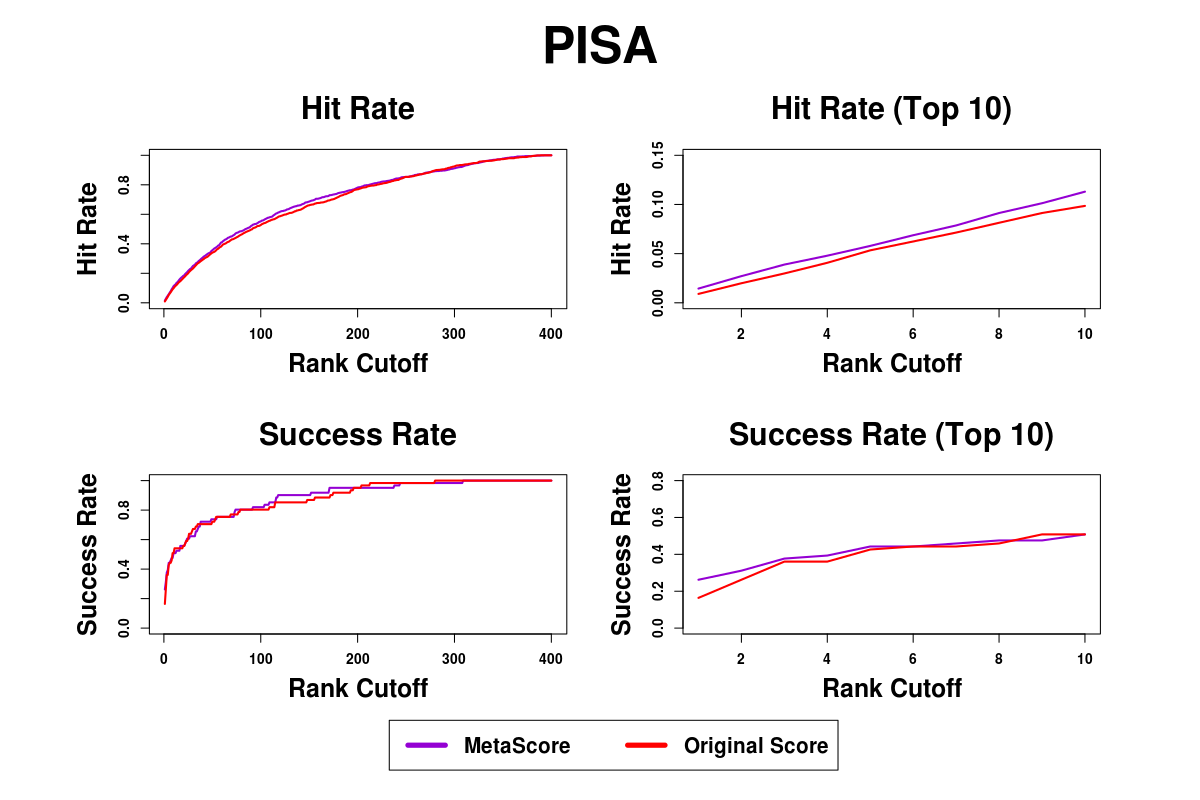

Supplementary Figure S7. Success rates and hit rates plotted against the top m conformations for a classical scoring method (PISA), machine learning-based method using RF, and the combined method of the two methods using BM4 decoy set. There are four panels. Top-left panel shows hit rates for conformations of top m ranging from 1 to 400; Top-right panel shows hit rates for conformations of top m ranging from 1 to 10; Bottom-left panel shows success rates for conformations of top m ranging from 1 to 400; Bottom-right panel shows success rates for conformations of top m ranging from 1 to 10.

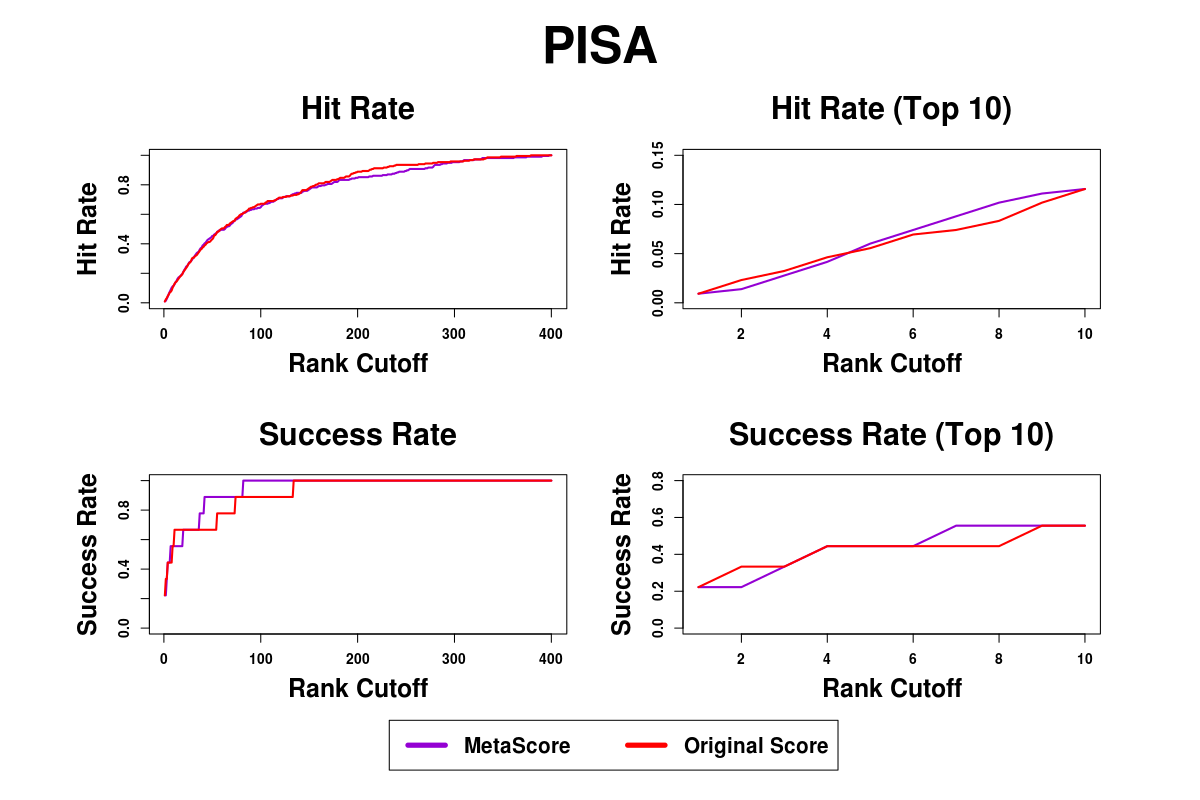

Supplementary Figure S8. Success rates and hit rates plotted against the top m conformations for a classical scoring method (PISA), machine learning-based method using RF, and the combined method of the two methods using BM5 decoy set. There are four panels. Top-left panel shows hit rates for conformations of top m ranging from 1 to 400; Top-right panel shows hit rates for conformations of top m ranging from 1 to 10; Bottom-left panel shows success rates for conformations of top m ranging from 1 to 400; Bottom-right panel shows success rates for conformations of top m ranging from 1 to 10.

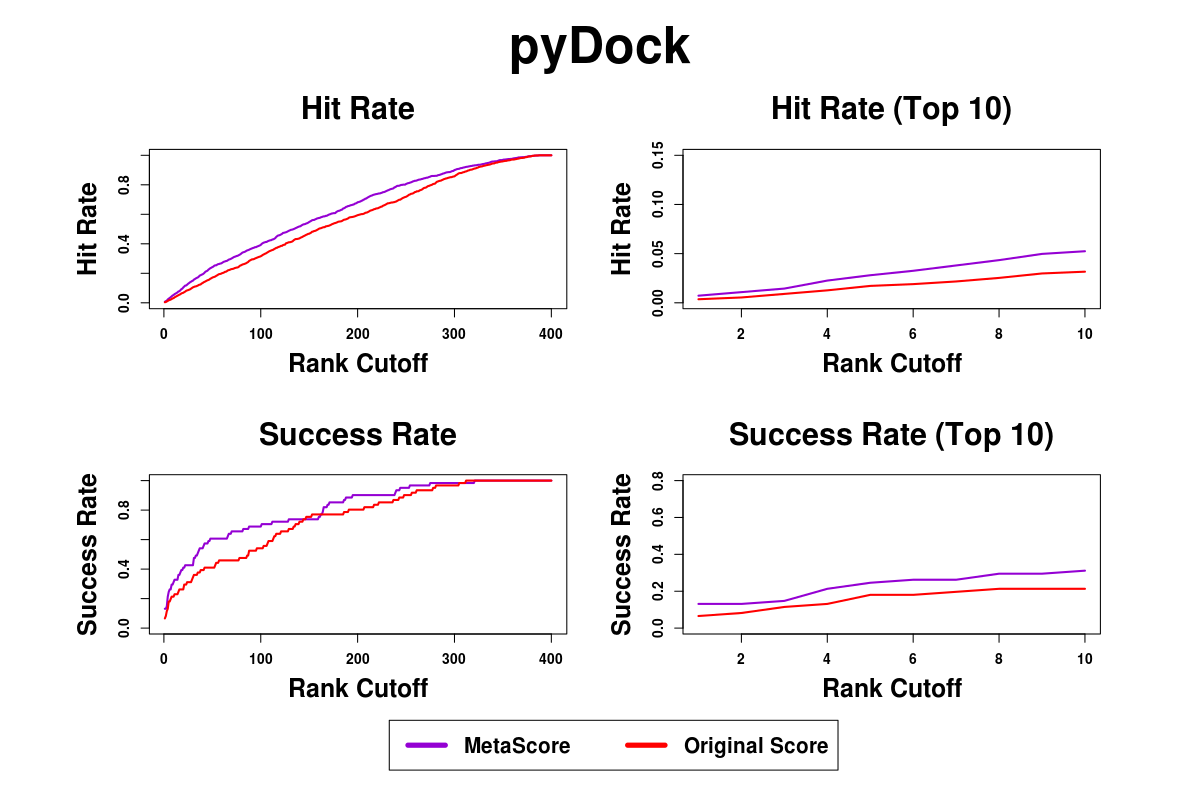

Supplementary Figure S9. Success rates and hit rates plotted against the top m conformations for a classical scoring method (pyDock), machine learning-based method using RF, and the combined method of the two methods using BM4 decoy set. There are four panels. Top-left panel shows hit rates for conformations of top m ranging from 1 to 400; Top-right panel shows hit rates for conformations of top m ranging from 1 to 10; Bottom-left panel shows success rates for conformations of top m ranging from 1 to 400; Bottom-right panel shows success rates for conformations of top m ranging from 1 to 10.

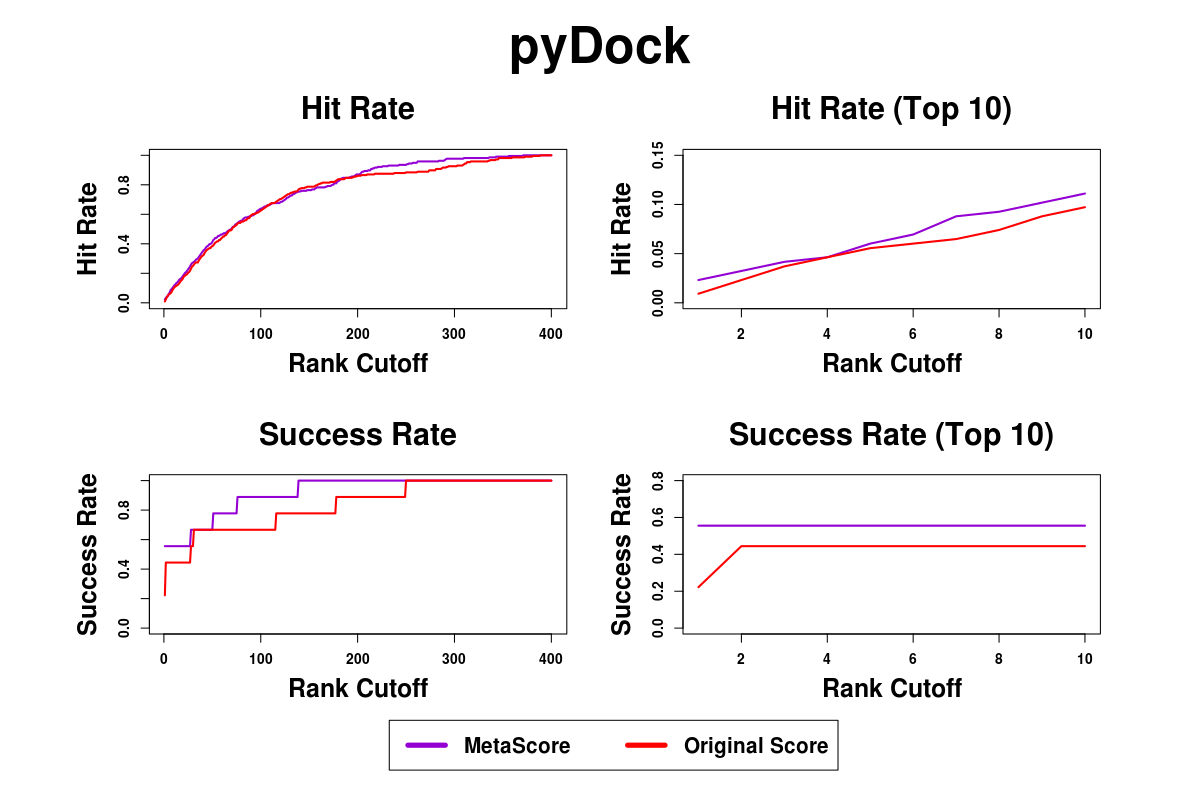

Supplementary Figure S10. Success rates and hit rates plotted against the top m conformations for a classical scoring method (pyDock), machine learning-based method using RF, and the combined method of the two methods using BM5 decoy set. There are four panels. Top-left panel shows hit rates for conformations of top m ranging from 1 to 400; Top-right panel shows hit rates for conformations of top m ranging from 1 to 10; Bottom-left panel shows success rates for conformations of top m ranging from 1 to 400; Bottom-right panel shows success rates for conformations of top m ranging from 1 to 10.

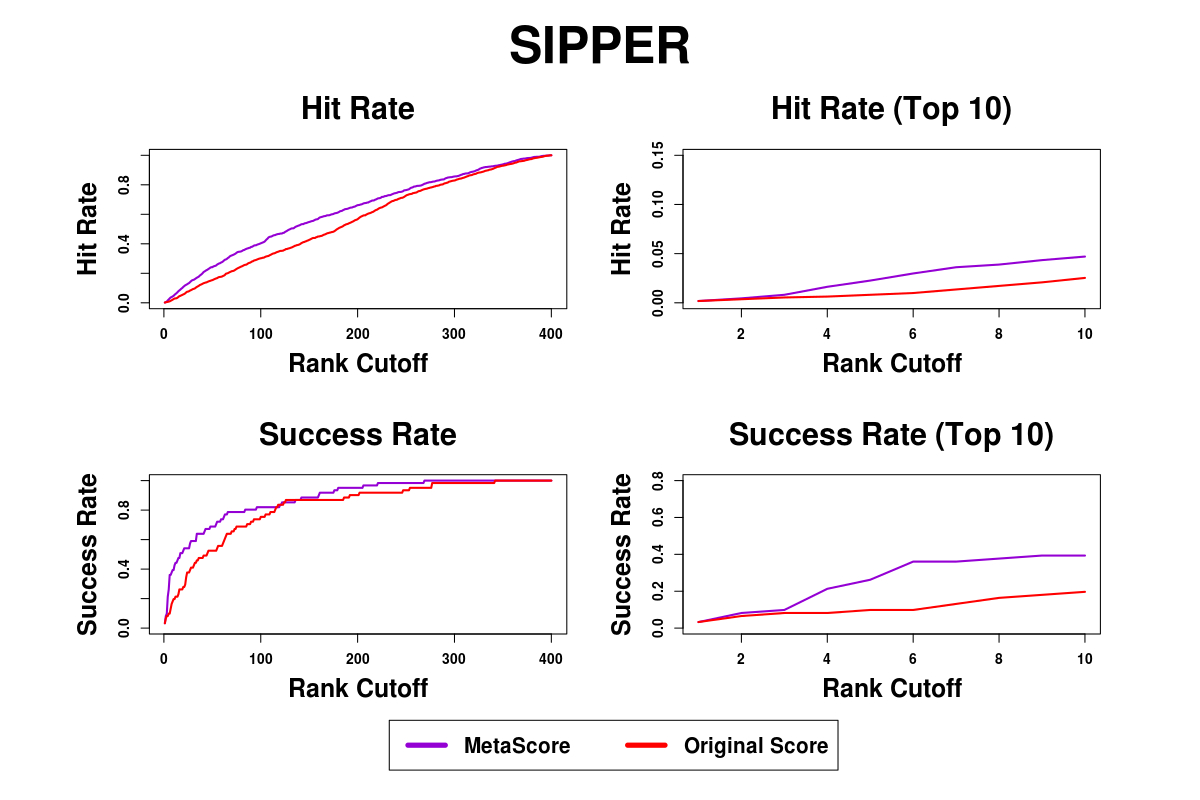

Supplementary Figure S11. Success rates and hit rates plotted against the top m conformations for a classical scoring method (SIPPER), machine learning-based method using RF, and the combined method of the two methods using BM4 decoy set. There are four panels. Top-left panel shows hit rates for conformations of top m ranging from 1 to 400; Top-right panel shows hit rates for conformations of top m ranging from 1 to 10; Bottom-left panel shows success rates for conformations of top m ranging from 1 to 400; Bottom-right panel shows success rates for conformations of top m ranging from 1 to 10.

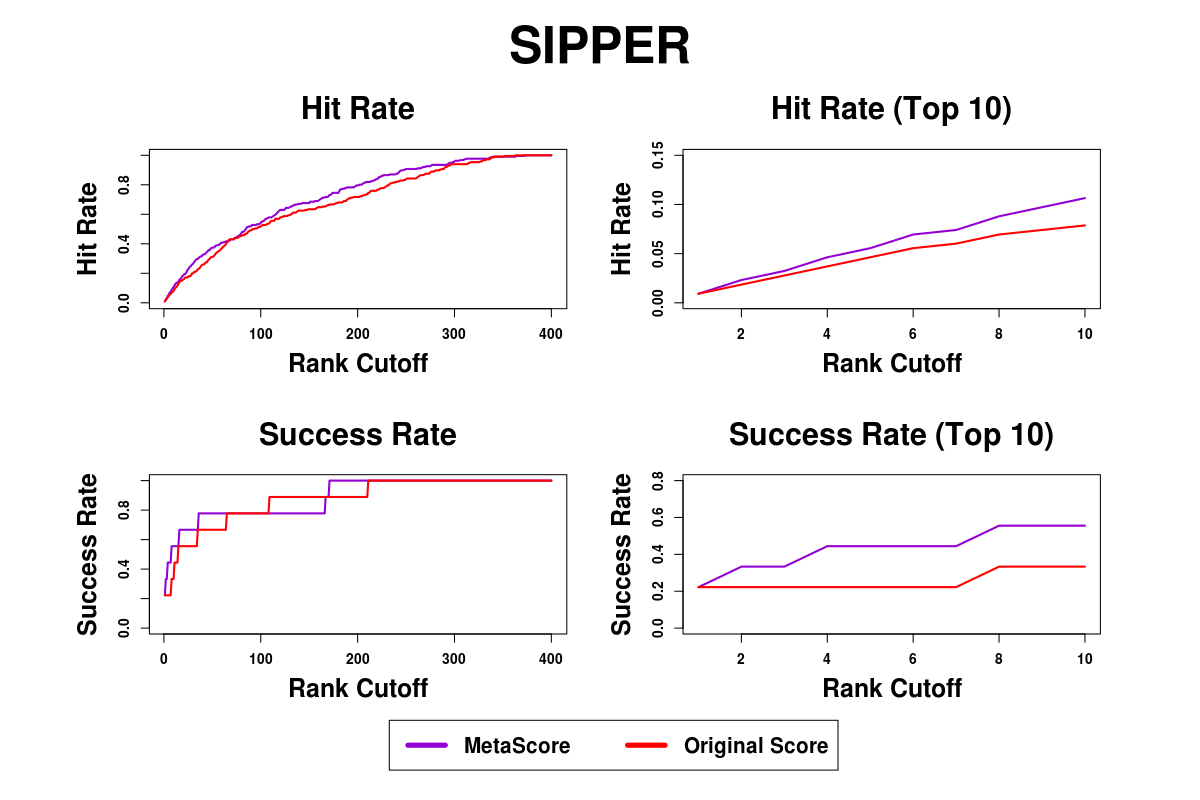

Supplementary Figure S12. Success rates and hit rates plotted against the top m conformations for a classical scoring method (SIPPER), machine learning-based method using RF, and the combined method of the two methods using BM5 decoy set. There are four panels. Top-left panel shows hit rates for conformations of top m ranging from 1 to 400; Top-right panel shows hit rates for conformations of top m ranging from 1 to 10; Bottom-left panel shows success rates for conformations of top m ranging from 1 to 400; Bottom-right panel shows success rates for conformations of top m ranging from 1 to 10.

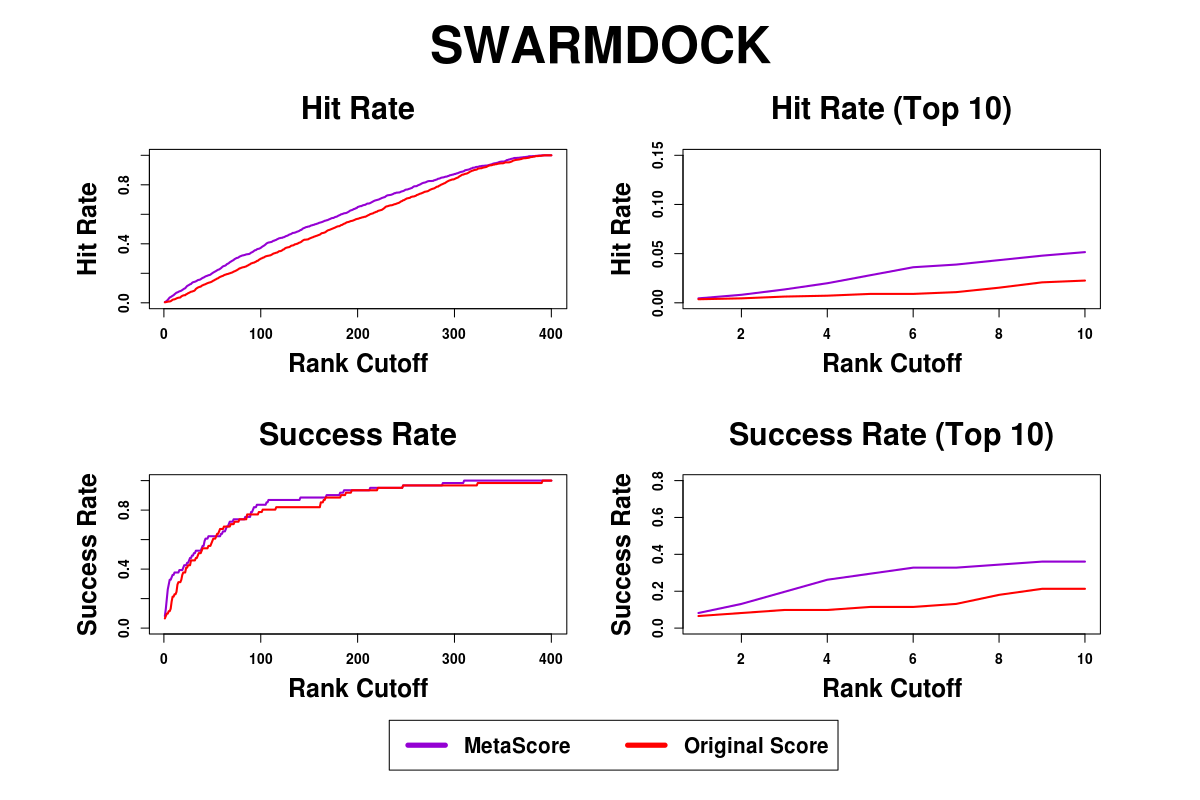

Supplementary Figure S13. Success rates and hit rates plotted against the top m conformations for a classical scoring method (SWARMDOCK), machine learning-based method using RF, and the combined method of the two methods using BM4 decoy set. There are four panels. Top-left panel shows hit rates for conformations of top m ranging from 1 to 400; Top-right panel shows hit rates for conformations of top m ranging from 1 to 10; Bottom-left panel shows success rates for conformations of top m ranging from 1 to 400; Bottom-right panel shows success rates for conformations of top m ranging from 1 to 10.

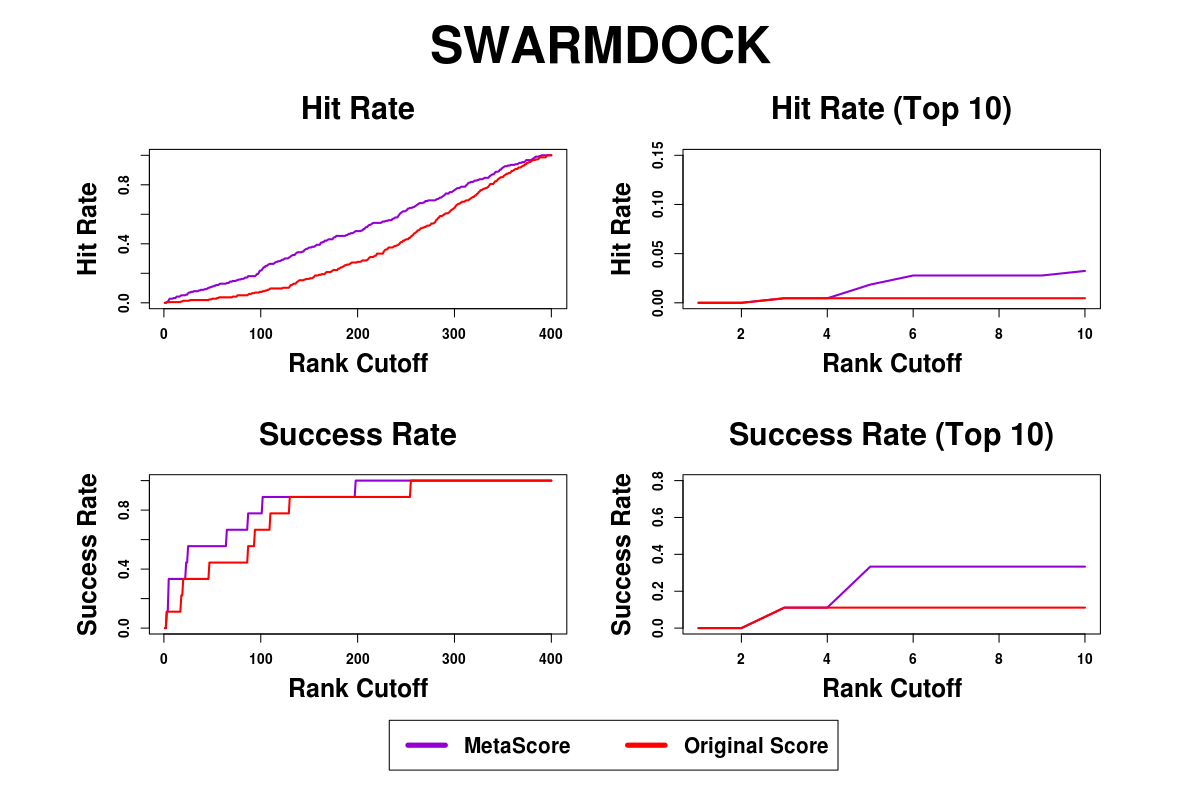

Supplementary Figure S14. Success rates and hit rates plotted against the top m conformations for a classical scoring method (SWARMDOCK), machine learning-based method using RF, and the combined method of the two methods using BM5 decoy set. There are four panels. Top-left panel shows hit rates for conformations of top m ranging from 1 to 400; Top-right panel shows hit rates for conformations of top m ranging from 1 to 10; Bottom-left panel shows success rates for conformations of top m ranging from 1 to 400; Bottom-right panel shows success rates for conformations of top m ranging from 1 to 10.

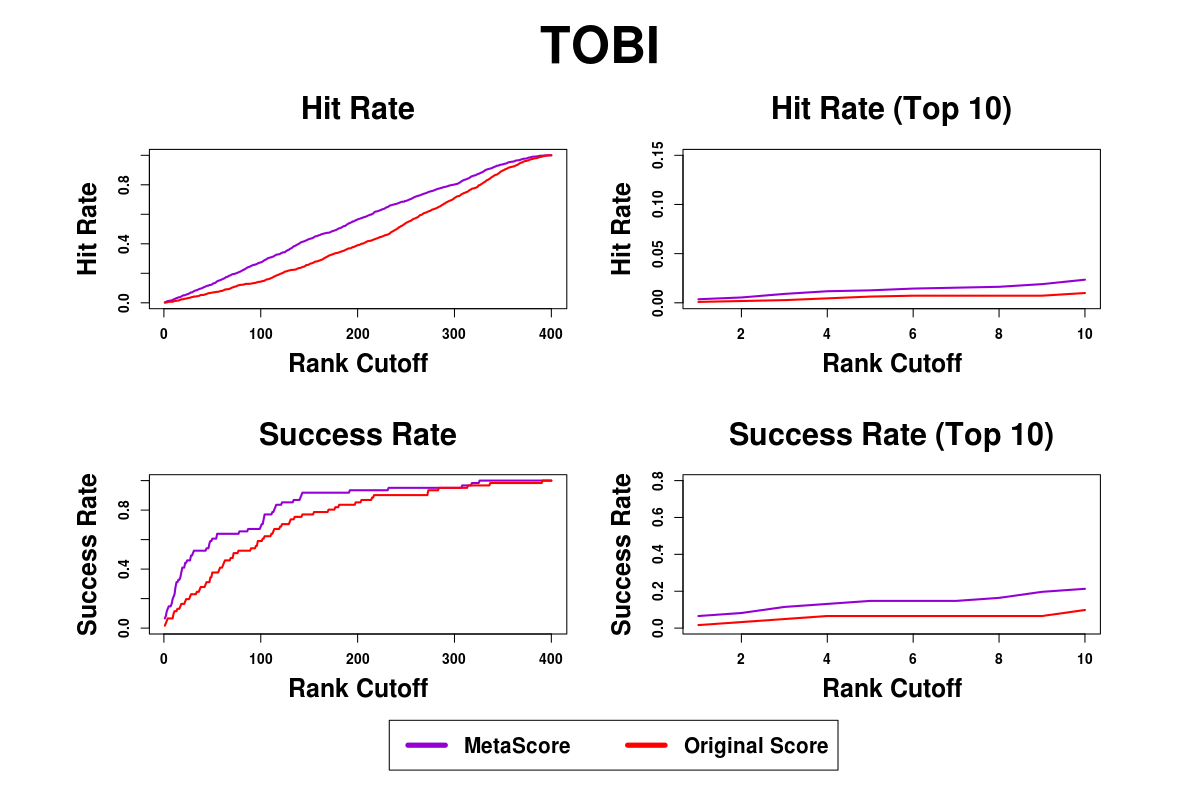

Supplementary Figure S15. Success rates and hit rates plotted against the top m conformations for a classical scoring method (TOBI), machine learning-based method using RF, and the combined method of the two methods using BM4 decoy set. There are four panels. Top-left panel shows hit rates for conformations of top m ranging from 1 to 400; Top-right panel shows hit rates for conformations of top m ranging from 1 to 10; Bottom-left panel shows success rates for conformations of top m ranging from 1 to 400; Bottom-right panel shows success rates for conformations of top m ranging from 1 to 10.

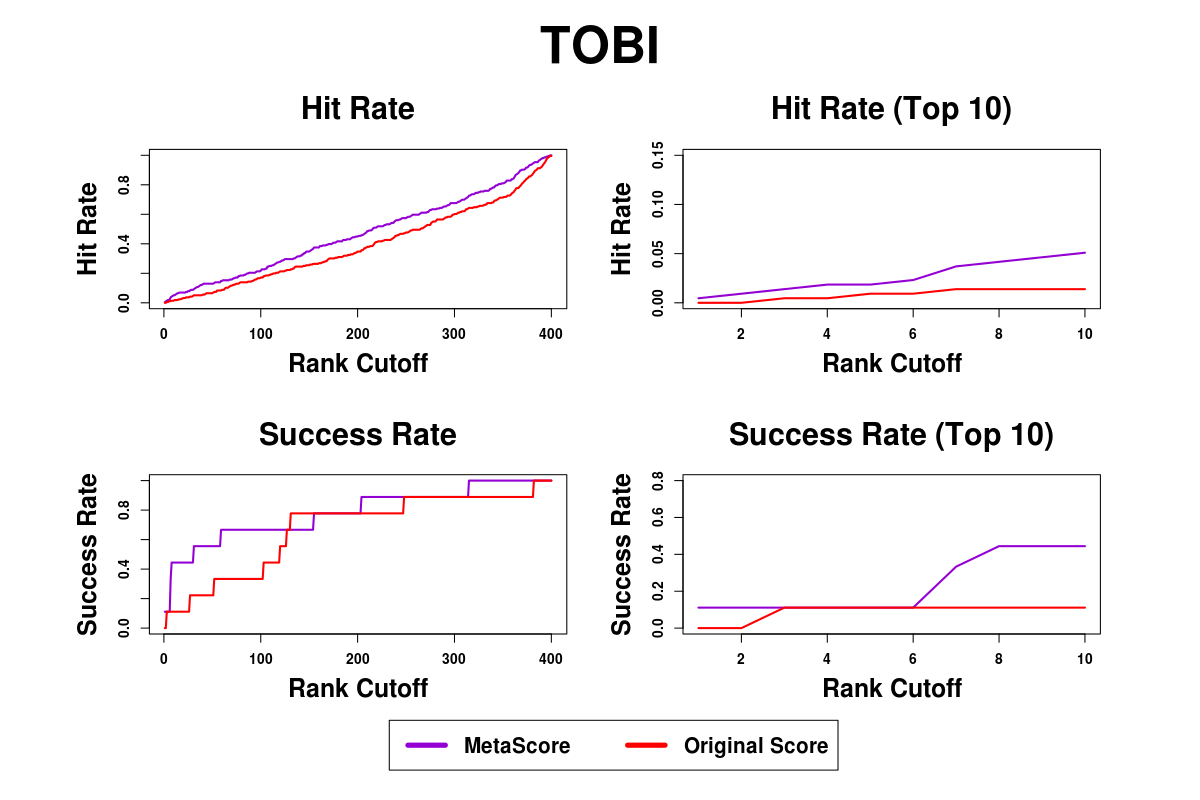

Supplementary Figure S16. Success rates and hit rates plotted against the top m conformations for a classical scoring method (TOBI), machine learning-based method using RF, and the combined method of the two methods using BM5 decoy set. There are four panels. Top-left panel shows hit rates for conformations of top m ranging from 1 to 400; Top-right panel shows hit rates for conformations of top m ranging from 1 to 10; Bottom-left panel shows success rates for conformations of top m ranging from 1 to 400; Bottom-right panel shows success rates for conformations of top m ranging from 1 to 10.

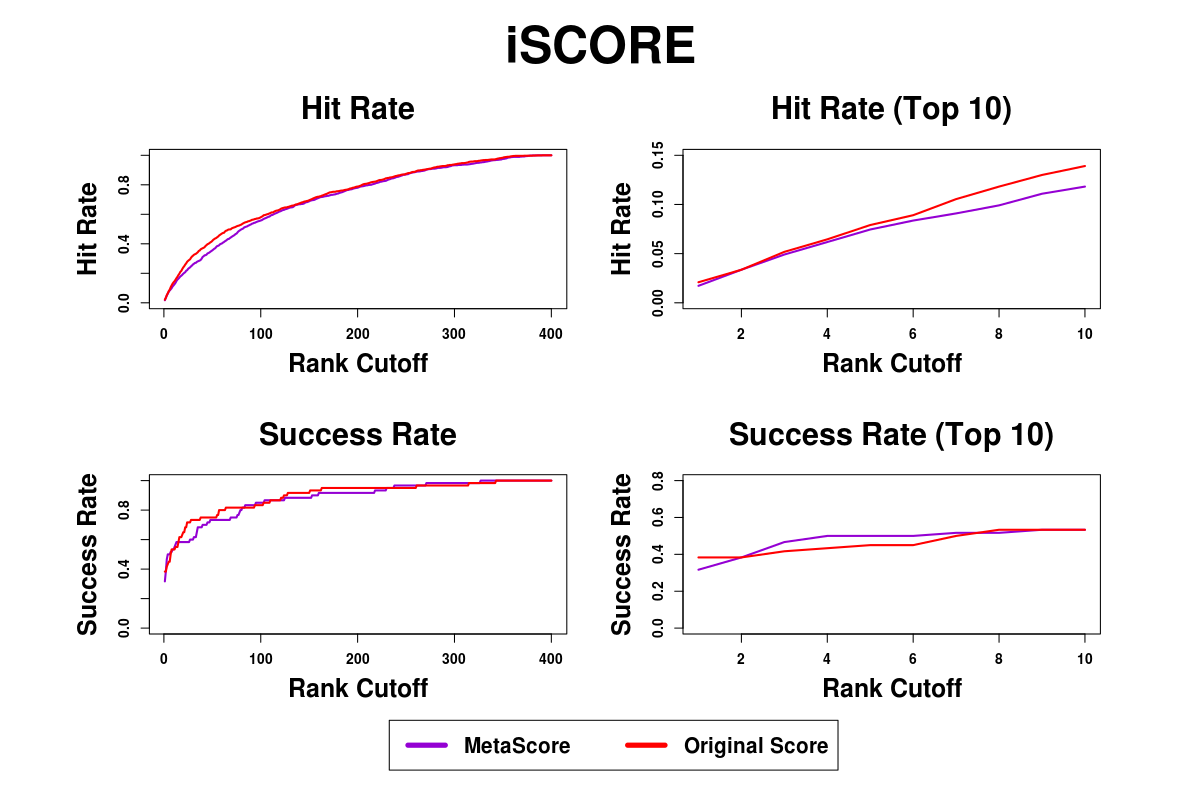

Supplementary Figure 17. Success rates and hit rates plotted against the top m conformations for a classical scoring method (iSCORE), machine learning-based method using RF, and the combined method of the two methods using BM4 decoy set. There are four panels. Top-left panel shows hit rates for conformations of top m ranging from 1 to 400; Top-right panel shows hit rates for conformations of top m ranging from 1 to 10; Bottom-left panel shows success rates for conformations of top m ranging from 1 to 400; Bottom-right panel shows success rates for conformations of top m ranging from 1 to 10.

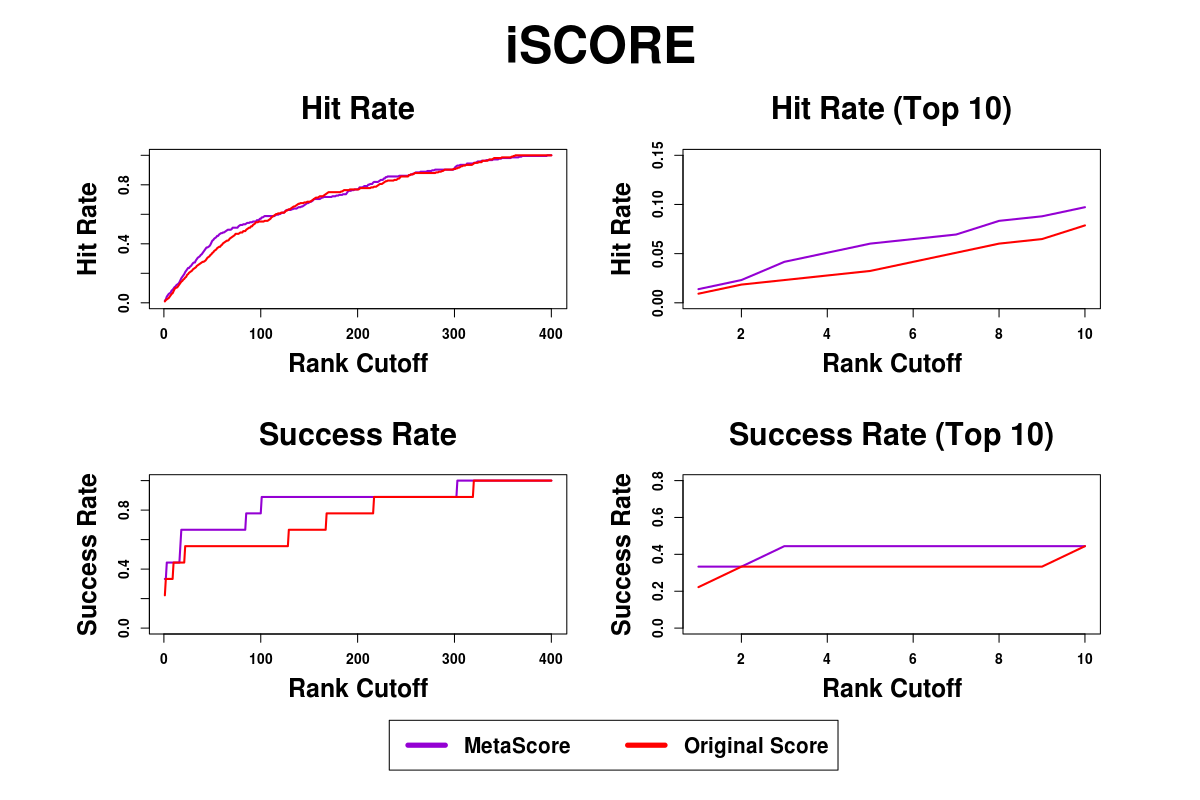

Supplementary Figure 18. Success rates and hit rates plotted against the top m conformations for a classical scoring method (iSCORE), machine learning-based method using RF, and the combined method of the two methods using BM5 decoy set. There are four panels. Top-left panel shows hit rates for conformations of top m ranging from 1 to 400; Top-right panel shows hit rates for conformations of top m ranging from 1 to 10; Bottom-left panel shows success rates for conformations of top m ranging from 1 to 400; Bottom-right panel shows success rates for conformations of top m ranging from 1 to 10.
